# Multi-policy models of interregional communication in the human connectome

**DOI:** 10.1101/2022.05.08.490752

**Authors:** Richard F. Betzel, Joshua Faskowitz, Bratislav Mišić, Olaf Sporns, Caio Seguin

## Abstract

Network models of communication, e.g. shortest paths, diffusion, navigation, have become useful tools for studying structure-function relationships in the brain. These models generate estimates of communication efficiency between all pairs of brain regions, which can then be linked to the correlation structure of recorded activity, i.e. functional connectivity (FC). At present, however, communication models have a number of limitations, including difficulty adjudicating between models and the absence of a generic framework for modeling multiple interacting communication policies at the regional level. Here, we present a framework that allows us to incorporate multiple region-specific policies and fit them to empirical estimates of FC. Briefly, we show that many communication policies, including shortest paths and greedy navigation, can be modeled as biased random walks, enabling these policies to be incorporated into the same multi-policy communication model alongside unbiased processes, e.g. diffusion. We show that these multi-policy models outperform existing communication measures while yielding neurobiologically interpretable regional preferences. Further, we show that these models explain the majority of variance in time-varying patterns of FC. Collectively, our framework represents an advance in network-based communication models and establishes a strong link between these patterns and FC. Our findings open up many new avenues for future inquiries and present a flexible framework for modeling anatomically-constrained communication.

## INTRODUCTION

Connectomes are network models of the brain’s structural connectivity (SC) and are fundamentally communication networks, shaping the flow of activity across the brain and facilitating communication events between distant neural elements [1–5]. The correlation structure of recorded brain activity – i.e. functional connectivity (FC) – reflects the time-averaged outcome of these processes [6].

While SC and FC can both be estimated from observation, e.g. non-invasively using functional and diffusion MRI, communication dynamics – the policies by which pairs of brain regions signal one another – are not directly observable [4]. One powerful approach for investigating communication policies is to simulate them *in silico*, and compare the outcome of those simulations with empirical observation [4, 7]. Most commonly, this means calculating a measure that denotes the ease of communication between pairs of brain regions and comparing these values with FC [8–20].

This framework has proven especially useful. While biophysical models offer biological plausible and dynamic perspectives on communication, they entail high computational costs and make specific assumptions about underlying biophysics that are not always empirically validated [21–24]. In contrast, communication models are computationally tractable (and are often analytically solvable) and generally do a better job explaining variation in FC [25].

However, there are a number of drawbacks to network communication models. First, adjudicating between models is, in general, not straightforward. Model fitness is typically defined *ad hoc* and the differences in performances between models – if model comparison is even performed – is often small [26]. Second, until recently [26, 27], communication models have been applied exclusively at the whole-brain level, i.e. the modeling framework assumes that every pair of brain regions communicates using the same policy. Finally, the incorporation of multiple competing policies within the same model has proven elusive. At present, the “state-of-the-art” is to generate non-interacting communication measures corresponding to different policies and combine them as explanatory terms in a multi-linear model [25, 28].

Here, we address these limitations directly by proposing a framework for modeling multiple, region-defined communication policies within the same framework. The policies we model exist along a continuum. On one extreme are decentralized and unbiased communication policies, e.g. random walks. On the opposite extreme are centralized and biased policies, e.g. shortest paths routing, in which knowledge about the network’s global topology is required to deliver a signal along the shortest path from a source to a target. Using this approach, we show that communication models in which policies are combined can outperform existing, single-policy models. Specifically, the best-performing models combine unbiased diffusive processes with directed and target-specific policies. We show that brain regions in the high-performing models all exhibit similar preferences – regions that comprise higher-order association systems favor communication by directed policies, while unimodal sensorimotor systems favor unbiased diffusion. Next, we apply the same framework to explain dynamic co-fluctuation patterns. We show that joint communication models explain, in some cases, nearly 60% of the variance in whole-brain network states. Finally, we show that preferences for unbiased, diffusive policies support the transient co-fluctuation between pairs of regions. The general modeling framework is flexible and can be easily extended to include triplets or quartets of communication policies, including policies not explicitly studied here.

## RESULTS

### Biased random walks for jointly modeling pairs of communication policies

Communication models aim to uncover the policies used by the brain to deliver signals from source to target regions. Under a given policy, it is often possible to calculate a measure of communication efficiency – the ease with which the signal gets delivered. For example, if the brain were to use shortest paths for communication, then path length could serve as a measure of communication efficiency. Presently, it is not possible to jointly model multiple interacting policies. In this and the next section, we outline a procedure for doing so and compare the fitness of joint communication models with traditional, single-policy models.

Briefly, our approach involves mapping commonly used communication policies onto biased and target-specific random walks. This allows us to parametrically combine multiple policies into a joint random walk in which the probability of a signal being delivered from one node to another is governed by two interacting policies. As an example, consider diffusion and shortest paths communication policies (Fig. 1). Diffusion can be modeled as an unbiased random walk in which a particle on a source node selects, at random, one of the source’s outgoing connections and hops to another node. This process repeats itself until, after a certain number of steps, the particle reaches its intended target (Fig. 1b).

**FIG. 1.**
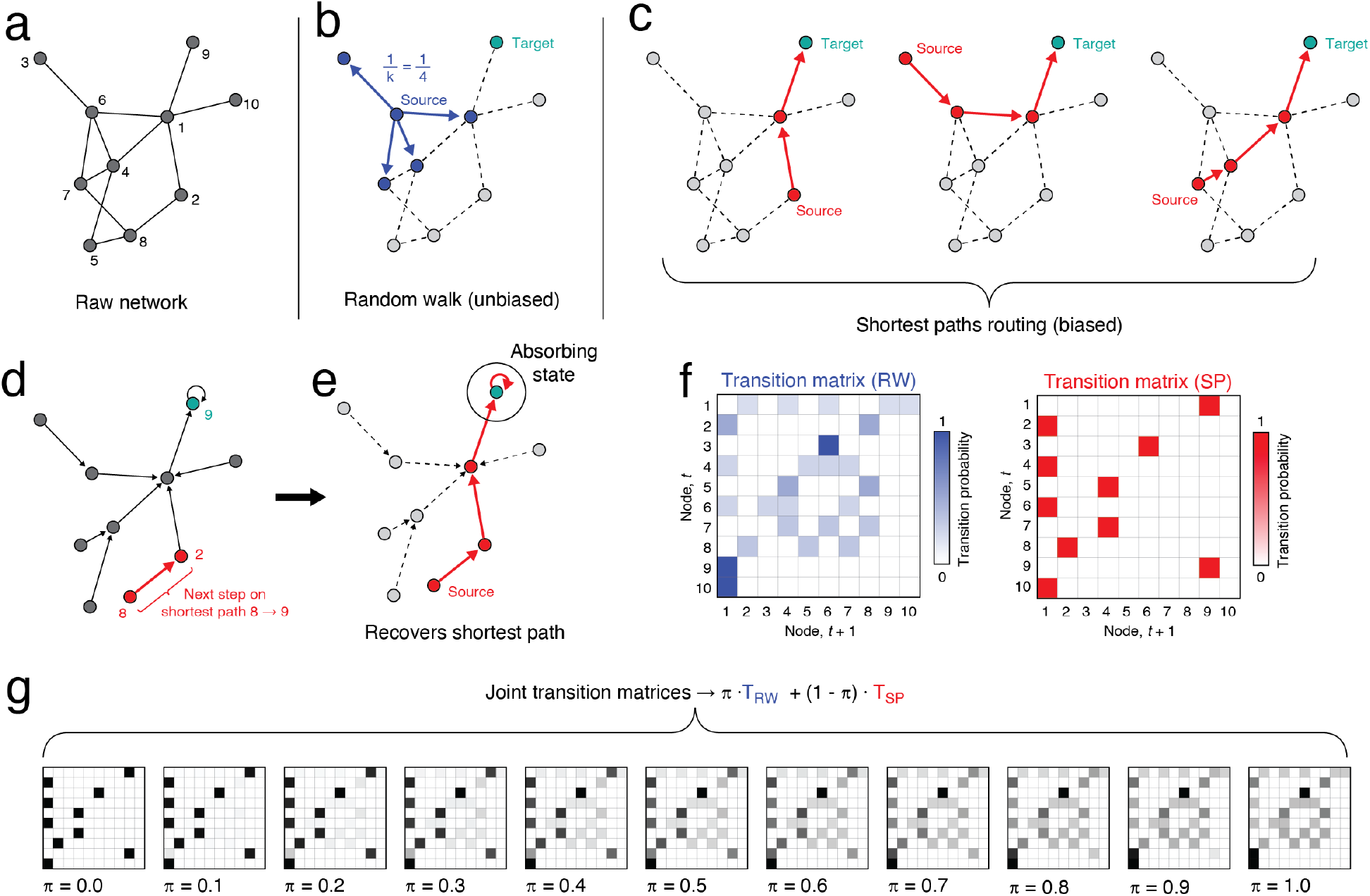
Mapping shortest paths routing onto random walks. Schematic illustrating how to map shortest paths routing models of communication onto a target-dependent random walk process. (*a*) Example network. (*b*) In an unbiased random walk, a particle “hops” from one node to another following outgoing connections. (*c*) In contrast, delivering a particle to the target node *via* shortest paths requires global knowledge of the network. Here, we show three example shortest paths (for a binary network). (*d*) For a given target node, however, we can construct a reduced network of directed connections so that, starting with any of the nodes in the network, a random walk would evolve towards the correct target. This is accomplished by retaining for every node a single outgoing connection (to the next node on the shortest path to the target). Note that this transformation of the network’s topology is dynamically equivalent to introducing a target-specific bias so that, under random walk dynamics, certain transitions are rendered impossible. (*e*) To ensure that once the node reaches the target it never leaves, we set the target node to be an absorbing state, i.e. has a self-connection. (*f*) Thus, we can construct two transition matrices – one describing a random walk over the full network and another describing a random walk over the reduced network. Given these two matrices, we can model a joint random walk process and construct a combined transition matrix as the linear combination of the these two under the condition that the sum of the weights is unity. (*g*) Examples of the joint transition matrix.

In contrast, a particle delivered using shortest paths routing selectively follows one route through the network, proceeding to its target in the fewest possible steps (Fig. 1c). However, it is possible to construct a reduced network by removing connections from the original, intact network such that, if one were to simulate a random walk on the network, the shortest paths to a target node are always accessed (Fig. 1d). This is accomplished by forcing node, *i*, to have a single outgoing connection that leads to the next node on the shortest path from *i* to the target node, *τ*, which we treat as an absorbing state (its outgoing connection is redirected back to itself; Fig. 1e). Effectively, the reduced network is equivalent to realizing a biased random walk on the original network, in which the only traversable edges are those that participate in shortest paths to the target.

Both the unbiased and biased random walks can be summarized by their transitions matrices, whose rows sum to unity and whose entries give the probability of transitioning from a node at step *t* to any other node at step *t* + 1 (Fig. 1f). To combine these two policies, we generate a joint transition matrix as a weighted combination of both. Provided that the weight, *π*, is bounded to the interval [0, 1], its specific value can be modulated to emphasize one or the other policy, leading to a continuum of possible random walks (Fig. 1g).

Note that when we generate transition matrices for the shortest paths policy (or any other target-dependent process), the corresponding transition matrix also varies as a function of target node. That is, because the shortest paths are variable across possible targets, we must generate a different biased transition matrix for each case (Fig. 2a). In contrast, the transition matrix for the unbiased random walk is identical, irrespective of target (Fig. 2b). Here, rather than generate a joint transition matrix using a globally defined weight [11], we introduce regional weights or “preferences” (Fig. 2a). Intuitively, the preference of region *i*, denoted as *π*_*i*_ ∈ [0, 1] indicates the likelihood that, should a particle arrive at node *i*, this node elects to use policy A (with probability *π*_*i*_) or policy B (with probability 1 −*π*_*i*_). We use the same set of preferences across all targets.

**FIG. 2.**
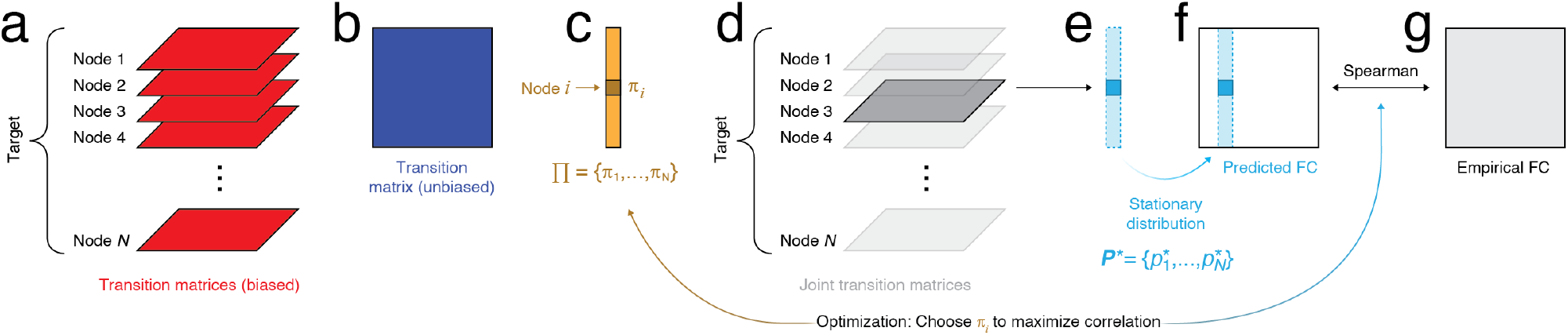
Linking joint communication models to FC. Here, we use the joint communication models to explain variance in FC. We do so by fitting region-specific preference parameters to pairs of communication processes. Here, we show an example using shortest paths routing and unbiased random walks as the two processes. (*a*) For the shortest paths process, we construct a reduced network and corresponding transition matrix for each possible target node. (*b*) In parallel, we construct a single transition matrix for the unbiased random walk using the full network. (*c*) To construct a joint transition matrix, we allow each node, *i*, to have its own preference for using random walks, *π*_*i*_∈ [0, 1]. The corresponding preference for the shortest path is 1−*π*_*i*_. (*d*). Consider a single target node, the set of preference parameters, and the corresponding joint transition matrices. (*e*) For each node, we can estimate the stationary distribution under this random walk, i.e. the expected configuration of random walkers over the network as the number of steps gets sufficiently large. (*f*) Repeating this process for every target node yields a different stationary distribution. The complete set of distribution vectors can be assembled into a *N*×*N* matrix. (*g*) We can then compare the elements of this matrix with that of the empirical FC matrix, e.g. as a Spearman correlation. We use this correlation magnitude as an objective function that we seek to optimize by selecting the appropriate values for Π = {*π*_1_, …, *π*_*N*_ }.

For a given target and set of preferences (Fig. 2d), we can calculate the stationary distribution of particles (Fig. 2e). That is, the probabilistic concentration of particles on each node as *t*→ ∞. If we repeat this process for all targets, we generate a matrix of stationary distributions (Fig. 2f). Like traditional single-policy communication models, the elements of this matrix can be linked statistically to the elements of the empirical FC matrix (Fig. 2g). Importantly, we also have the ability to modify the regional preferences so as to strengthen the statistical correspondence between the matrix of stationary distributions and FC.

### Comparing joint communication models with traditional communication measures

In the previous section we described a framework for jointly modeling multiple communication policies simultaneously. Here, we systematically evaluate the performance of multi-policy models and compare it against 46 parameterizations of 10 distinct communication measures that appear in the extant literature.

As part of this approach, we pursue two distinct analysis pipelines, both aimed at linking structural and functional connectivity (Fig. 3a). First, following [26], we generate fully-weighted predictor matrices corresponding to 46 parameterizations of 10 distinct communication policies that. For each predictor, we calculated the correlation (Spearman) of its elements with those in the static FC matrix (Fig. 3b). In effect, this first approach tests existing methods and establishes a clear baseline for performance.

**FIG. 3.**
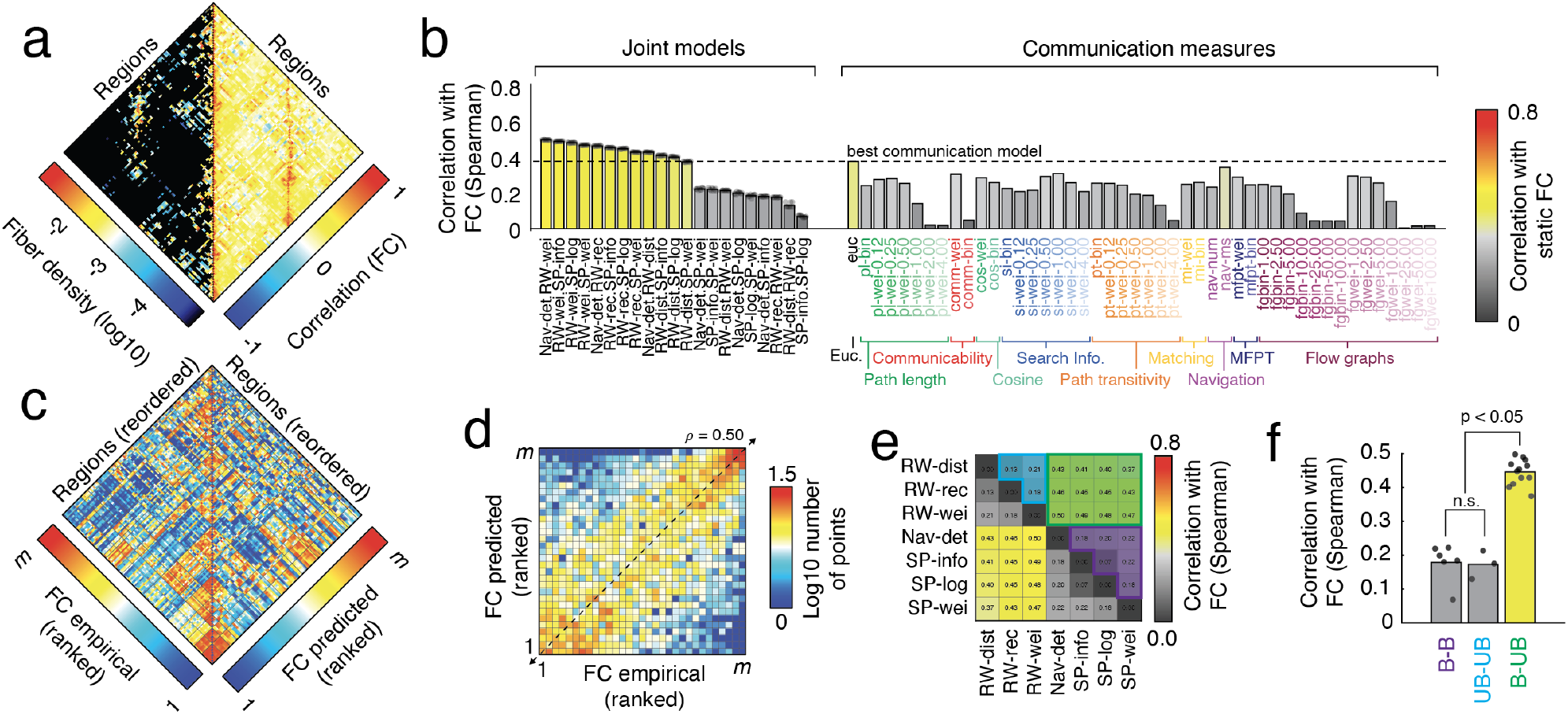
Connectivity matrices and comparison of joint models with communication measures. (*a*) We analyzed structural and functional connectivity (SC; FC) data at the group level. The SC matrix (left) was sparse; existing edge weights corresponded to fiber densities. The FC matrix (right) was fully weighted and signed; edge weights correspond to interregional correlations. (*b*) We fit regional preference values for 21 joint models (combinations of two communication policies) by optimizing the correspondence (Spearman rank correlation) between the stationary distributions of random walkers and the FC matrix (points represent model fitness over multiple repetitions of the optimization algorithm using different initial conditions). In parallel, we calculated 46 communication measures based on 10 parameterized families of policies. (*c*) Rank-transformed empirical FC (left) and FC predicted by the best two-policy communication model. (*d*) Two-dimensional histogram depicting the correlation of empirical and predicted FC following the rank transformation. (*e*) Mean fitness of the 21 joint-policy models ordered into 7×7 matrix (rows and columns correspond to individual communication policies). Purple, blue, and green blocks correspond to joint models that pair shortest path policies with other shortest path policies, random walk policies with other random walk policies, and shortest path with random walk policies. Note that we group the navigation policy with the shortest paths policies as both are target-dependent, i.e. the transition matrix varies depending on target node. (*f*) The upper triangle elements of the performance matrix grouped by model category.

In parallel, we used the framework described in the previous section to model 21 multi-policy processes (based on seven individual policies). In general, these policies came from one of three families: unbiased random walks, shortest paths routing, and greedy navigation. More specifically, we considered the following set of seven policies (and their 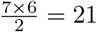 combinations):

1. `SP.wei`: we calculate the shortest path from node *i* to the target, *τ*, based on a reciprocal transformation of weight to cost, i.e.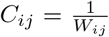. We then write the biased random walk as *T*_*ij*_ = 1 if *j* is the next node on the shortest path from *i* to *τ* and 0 otherwise.
2. `SP.log`: we calculate the shortest path from node *i* to the target, *τ*, based on a log transformation of weight to cost, i.e.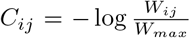. We then write the biased random walk as *T*_*ij*_ = 1 if *j* is the next node on the shortest path from *i* to *τ* and 0 otherwise.
3. `SP.info`: we calculate the shortest path from node *i* to the target, *τ*, based on a log transformation of weight to information theoretic cost in bits, i.e. 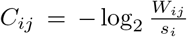, where *s*_*i*_ = Σ_*j*_*W*_*ij*_ is the weighted degree of node *i*. We then write the biased random walk as *T*_*ij*_ = 1 if *j* is the next node on the shortest path from *i* to *τ* and 0 otherwise.
4. `Nav.det`: navigation is a decentralized and greedy heuristic for delivering a particle from node *i* to a target node *τ* with spatial co-ordinates {*x*_*τ*_, *y*_*τ*_, *z*_*τ*_ }. Specifically, this policy considers the neighbors of *i*, Γ_*i*_ = {*j*, …, *k*} and their respective straight-line distances from the target, Δ_*i*_ = {*d*_*jτ*_, …, *d*_*kτ*_ }, where 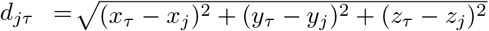. The particle is then delivered to the node nearest the target.
5. `RW.wei`: an unbiased random walk over the network in which transition probability is given by, 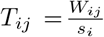.
6. `RW.dist`: an unbiased random walk over the network in which transition probability is given by, 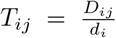, where *D*_*ij*_ is the Euclidean distance of the connection between regions *i* and *j* and *d*_*i*_ = Σ _*j*_ *D*_*ij*_ is a normalization factor equal to the total length of all connections incident upon node *i*. If no connection exists then *D*_*ij*_ = 0. Note that this random walk leads to a preference for the walker to select long range connections.
7. `RW.rec`: an unbiased random walk over the network in which transition probability is given by,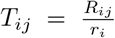, where *R*_*ij*_ is the correlation similarity 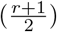 between neurotransmitter receptor profiles of regions *i* and *j* with entries zeroed out if no structural connection exists [29]. The denominator *r*_*i*_ = Σ _*j*_ *R*_*ij*_ is the weighted degree of this matrix.

These seven policies can be divided, roughly into two groups. `SP.wei, SP.log, SP.info`, and `Nav.det` represent target-specific policies and, in order to be modeled as a Markov chain, require that we bias the transition probabilities. For this reason, we refer to this subset of policies as the “biased” group. The remaining policies, `RW.wei, RW.dist`, and `RW.rec`, are fully decentralized, target-agnostic, and require no biases to be modeled as Markov Chains. For this reason, we refer to this second subset of policies as “unbiased”.

For every unique pair of policies, we fit local preferences, Π ={*π*_1_, …, *π*_*n*_}, so as to maximize the Spearman correlation between the stationary distribution of walkers and the static FC. Note that while other studies have focused on predicting FC for only mono-synaptically coupled pairs of brain regions [6], we predict FC for all pairs of brain regions irrespective of whether their connections are mono- or poly-synaptic. This is the case for both the traditional communication models as well as the multi-policy models.

We found that, of the traditional communication models, Euclidean distance (*ρ* = 0.38), navigation (*ρ* = 0.34), search information (*ρ* = 0.31), and communicability (*ρ* = 0.31) were among the top performers (Fig. 3b). These results are broadly in line with other recent studies [9, 10, 26].

In comparison, we found that, of the 21 fitted multi-policy models, eleven (52.4%) outperformed the best communication model. The top performers included `Nav.det-RW.wei` (*ρ* = 0.50), `SP.info-RW.wei` (*ρ* = 0.49), `SP.log-RW.wei` (*ρ* = 0.48), and `SP.wei-RW.wei` (*ρ* = 0.47). As an example, we show the empirical and predicted FC for the top-ranked model in Fig. 3c,d.

Notably, we also compared the observed fitness values against those obtained under two network null models, i.e. identical model-fitting procedures but with randomized SC. Both null models exactly preserved nodes’ degrees and approximately preserved nodes’ strengths. Additionally, one of the models also approximately preserved the total wiring cost of the network. In general, we found that high-performing models predicted static FC above and beyond both null models (*p <* 0.01; Fig. S1).

Collectively, these results suggest that multi-policy and regionally-heterogeneous communication models outperform existing measures in terms of their ability to predict the organization of FC. This improvement in performance is reduced when the underlying anatomical connectivity is perturbed by random and space-constrained rewiring algorithms. Nonetheless, our findings suggest that multi-policy models do not, in all cases, outperform existing measures; many combinations of policies lead to performance that is comparable to or worse than traditional communication models, suggesting that the specificity and synergy among policies is largely responsible for the improvement.

### Pairing decentralized and target-specific policies yields improved fit to static FC

In the previous section, we demonstrated that joint models yielded stronger structure-function correlations than traditional communication measures. In this section, we focus specifically on the joint models and seek to identify principles that explain the heterogeneity in model performance.

Of the 21 joint models, we divided them into three categories: those that combine two unbiased diffusive processes, those that combine two biased, target-dependent processes, and those that combine unbiased and biased policies in the same model (Fig. 3e). Interestingly, we find that combining biased and unbiased policies yield improved performance relative to the other categories (t-test, *t*(19) = 14.15, *p* = 1.5 ×10^−11^; Fig. 3f).

These results suggest that, although the joint models include more parameters than traditional communication models, there is considerable heterogeneity in terms of performance from one model to another. In fact, many of the multi-policy models perform poorly relative to traditional communication models. These observations suggest that the complexity of the fitted multi-policy policies alone does not fully explain why subsets of these models as well as they do. Further, our findings suggest that much of this heterogeneity can be parsimoniously explained based on the kind of communication policies paired together in the same model. Finally, these observations motivate exploring the spatial structure of the regional preferences for one communication policy *versus* another.

### Regional preferences divide along task positive/negative divisions of cerebral cortex

To this point, we have demonstrated that joint models outperform traditional communication measures in terms of explaining variance in FC. Further, we showed that the best-fitting models tended to pair unbiased diffusion policies with target-dependent policies. However, we have not explicitly examined the regional preferences for one policy or the other, i.e. the vectors Π = {*π*_1_, …, *π*_*N*_}. In this section, we examine the stability of these patterns over multiple runs and characterize their spatial distribution across the cerebral cortex.

First, we assessed whether repeated runs of optimization algorithm yielded similar regional preferences. To do this, we repeated the optimization algorithm 10 times for each joint model and calculated the Spearman rank correlation between all 10 sets of preference vectors (Fig. 4a). In general, we found that models were highly dissociable from one another (mean similarity between runs of the same and different model were *μ*_*s*_ = 0.74±0.20 and *μ*_*d*_ = 0.25±0.25; t-test *t*(21943) = 57.5, *p <* 10^−15^; Fig. 4b). We also found that models combining unbiased and biased diffusion policies converged to more similar solutions than other models (*t*(943) = 31.3, *p <* 10^−15^).

**FIG. 4.**
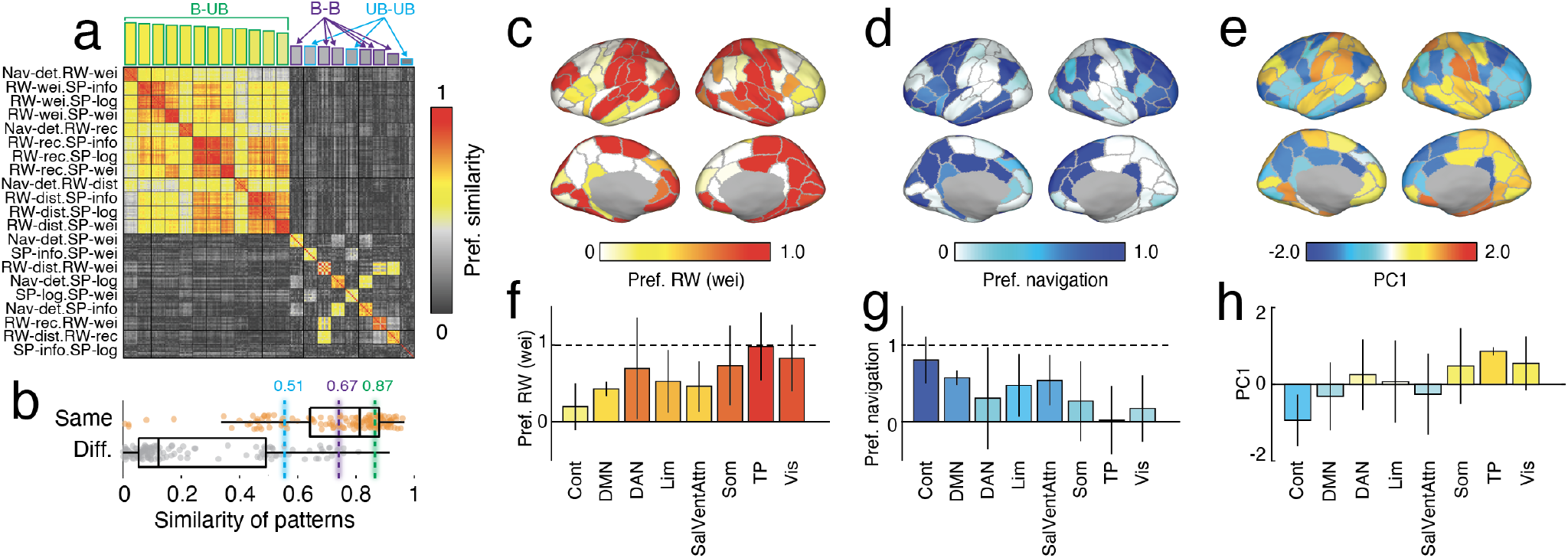
Stability and comparison of regional preference patterns. (*a*) We repeated the optimization algorithm ten times for every joint model, each time obtaining an estimate of regional preferences. (*a*) Here, we compare the similarity of those preference vectors (Spearman correlation) within repeated runs of the same model and between different models. (*b*) We partition models into three groups and show that joint models in which one of the policies is unbiased and diffusive and the other is biased and target-dependent converge to more stable solutions with less variability across successive runs. Panels *c, d, f*, and *g* regional preferences for a weight-based random walk (red colors) and navigation (blue colors) projected onto the cortical surface and aggregated by brain systems. Rather than focusing on a single model, we also aggregated regional preference vectors for all models that paired biased with unbiased policies and performed a principal components analysis on these patterns. Panel *e* shows the first principal component projected onto the brain surface and aggregated by brain systems.

Next, we examined the spatial patterning of the models. Initially, we focused on the top-performing model – `Nav.det-RW.wei` (Fig. 4c,d). We found that the optimal preferences were largely bilaterally symmetric, and were characterized by a pattern in which higher-order association areas, including control, default mode, and salience/ventral attention networks favored the navigation policy. In contrast, preferences for the unbiased, weight-based diffusion policy favored primary sensory (visual and somatomotor), dorsal attention, and temporo-parietal networks (Fig. 4f,g).

Finally, we noted that, although not identical, joint models combining unbiased and biased diffusion policies tended form a “block” (see the green block in the similarity matrix depicted in Fig. 4a), indicating that, although preferences were distinct for each model, all models of this type were mutually similar with one another. Accordingly, we pooled preferences from these models and performed a principal components analysis, yielding a first component (PC1) that explained 66.7% of variance. When we examined the spatial structure of this component, we found that it defined a mode of variation in which higher-order association cortices (and the control network in particular) favored the biased diffusion policies, whereas sensorimotor systems favored the unbiased policy (Fig. 4e,h), mirroring the results shown in Fig. 4c,d,f,g.

Importantly, we found that the regional preference values were not clearly linked to low-level features of structural connectivity. Specifically, we focused on regional degree and strength. The mean Spearman correlation of those measures with regional preferences were *ρ*_*strength*_ = 0.25 ± 0.15 and *ρ*_*degree*_ = 0.17 ± 0.10 (see Fig. S2).

Collectively, these results suggest high-performing models discover similar preference patterns. Specifically, to match static FC patterns, regions in sensorimotor and dorsal attention networks exhibit a preference for unbiased diffusive communication policies, while regions in higher-order association cortices, namely the control system, prefer biased and target-dependent policies.

### Linking joint communication policies to dynamic cofluctuations

To this point, we have focused on using joint communication models to predict static, time-averaged FC. However, there is increasing evidence that connectivity patterns estimated with fMRI BOLD fluctuate on timescale of 10’s of seconds. Here, we shift our target away from static FC to dynamic network states and assess whether joint communication models can predict spontaneous cofluctuation patterns.

To address this question, we leveraged a recently developed approach for tracking moment-to-moment changes in co-fluctuations [30, 31]. Briefly, this approach “unwraps” functional connections across time, yielding a framewise account of an edge’s magnitude. Previous studies using this approach demonstrated that the collective behavior of these “edge time series” across the entire cerebral cortex results in “events” – brief instances in time when many edges simultaneously exhibit high-amplitude co-fluctuations [30, 32]. Although short-lived and infrequent, the mean co-fluctuation over these brief periods of time closely approximates the pattern of static FC. More recently, it was shown that these can be divided into a number of approximately repeating clusters.

Here, we calculate edge time series for 70 individuals, estimated events on a per-subject basis, pool the event co-fluctuation patterns together, and, following [33, 34], apply a data-driven clustering algorithm to these matrices, yielding three large states. The first pattern is typified by the collective co-fluctuations of visual, somatomotor, and dorsal attention networks with one another. The second cluster involves co-fluctuations of control, default mode,and salience/ventral attention networks. Finally, the third cluster, which was more diffuse and less selective, involved co-fluctuations of all brain areas other than the visual network, but especially components in the salience ventral attention network with sub-components of the control, default mode, and somatomotor networks.

Following the same procedure we applied to static FC, we fit 21 joint models to predict the centroids for each of the three clusters. We made three important observations. First, we found that joint communication models provided better explanations of individual centroid co-fluctuation patterns than of static FC. The top model for states 1, 2, and 3 achieved correlations of *ρ* = 0.58, *ρ* = 0.67, and *ρ* = 0.77 (Fig. 5), respectively (in comparison, the best model for static FC achieved a correlation of *ρ* = 0.50). Second, we found that best-fitting model for the three states was dissimilar from the one that best-predicted static FC. Specifically, we found that the best-fitting state models for states 1, 2, and 3 were `RW.wei-SP.info`, `RW.rec-Nav.det`, and `RW.wei-SP.wei`; see Fig. 5g and Fig. 6a,b,f,g,k,l).

**FIG. 5.**
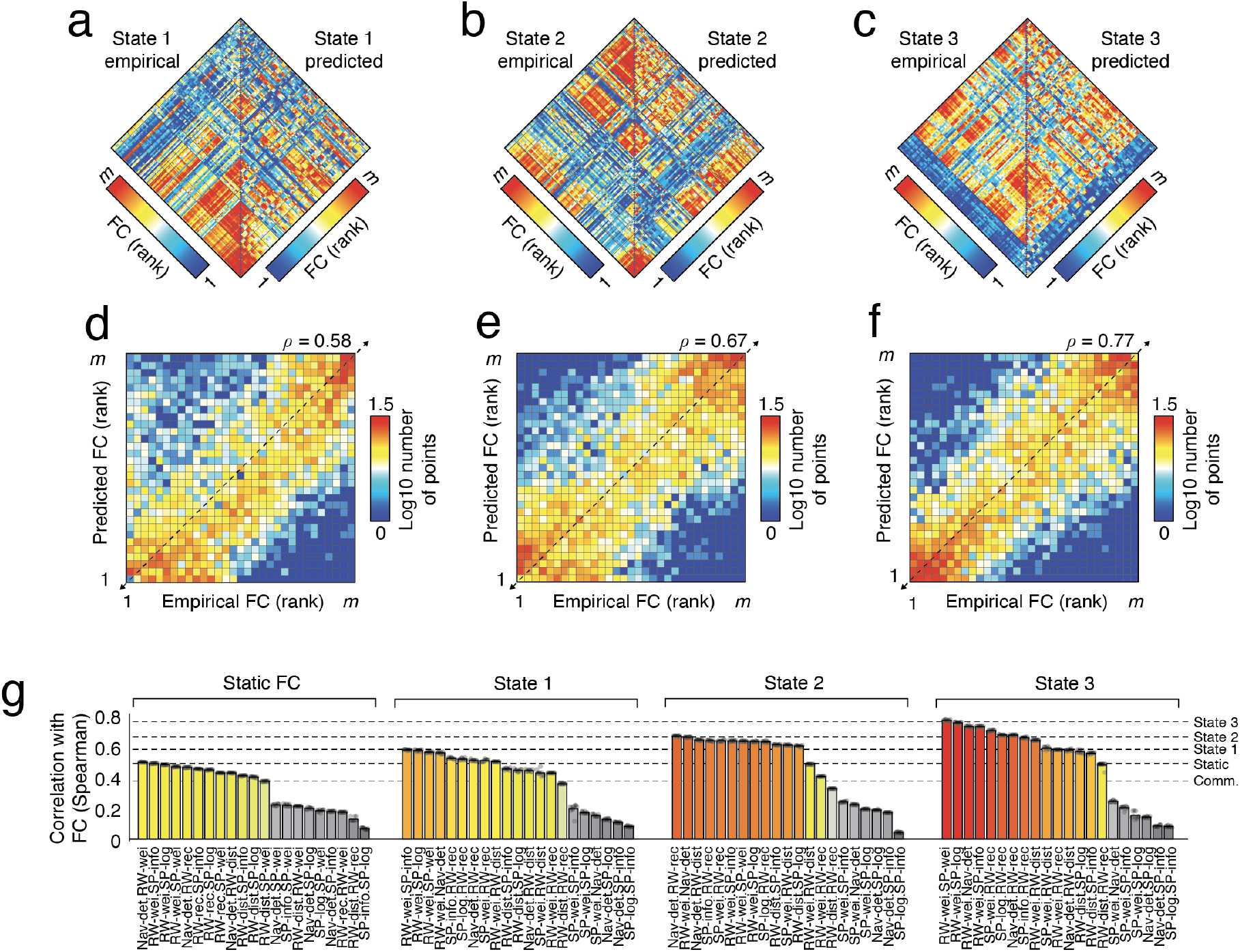
Predicting time-varying co-fluctuation patterns with joint communication policies. In addition to predicting static FC, we used joint communication models to predict co-fluctuation patterns estimated using “edge time series”. Panels *a*-*c* depict the co-fluctuation patterns for three states. On the left of each diamond plot is the empirical pattern estimated from data. On the right side of each plot is the predicted co-fluctuation by the model. Note that all connection weights have been transformed into ranks. Panels *d* -*f* show two-dimensional scatterplots of the rank-transformed empirical and predicted co-fluctuation values. (*g*) Comparison of model fitness for static FC and for states 1, 2, and 3 estimated from edge time series. Dashed lines represent the best-performing models along with the best-performing of the traditional communication models.

**FIG. 6.**
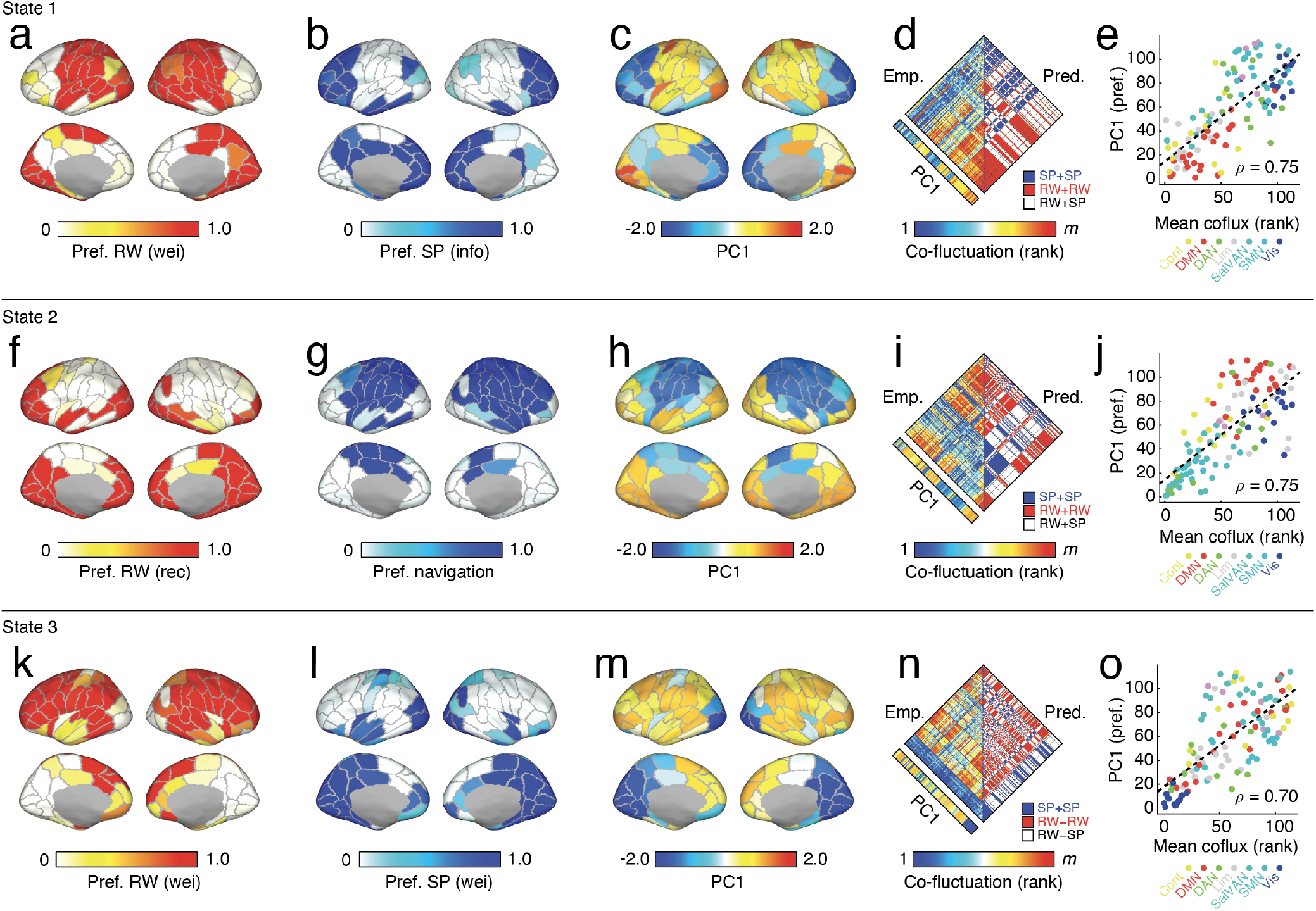
Regional preferences for time-varying co-fluctuation patterns. Each row in this figure depicts results associated with a different co-fluctuation “state”. Panels *a*-*e, f* -*j*, and *k* -*o* correspond to states 1, 2, and 3, respectively. (*a*) Regional preferences for the unbiased random walk term. (*b*) Regional preferences for the biased and target dependent policy. (*c*) We retained and concatenated preference vectors for high-performing joint policies (*ρ >* 0.4) and performed a principle components analysis on those vectors. Panel *c* depicts the first principal component. (*d*) The left-hand side of this diamond plot depicts the rank-based co-fluctuation pattern for state 1. On the right-hand side, we assign edges one of three possible labels. Red edges link nodes that both prefer the unbiased random walk. Blue edges link nodes that both prefer the biased, target-dependent random walk. White edges link nodes that prefer different policies. Note that the red edges on the right-hand side closely overlap with the strongest co-fluctuations in the cluster centroid map on the left. (*e*) Correlation of the first principal component (PC1) with the mean co-fluctuation of empirical state-based FC matrix. Here, we rank-transform each vector before plotting. Points are colored based on their system assignment.

Finally, we found that the preference vectors were also dissimilar from that of those associated with static FC and that the spatial structure of these preferences also varied across the three states (Fig. 6). As before, we also assessed whether preference patterns were similar across high-performing models (in this case, those with *ρ*≥0.4). Of those models, we calculated the mean preference pattern of all repetitions, concatenated these patterns into a region-by-model matrix, and performed a principal components analysis (the first PC accounting for 59%, 57%, and 59% of total variance across preferences for states 1, 2, and 3, respectively; see Fig. 6c,h,m for projections of these components onto the cortical surface). We show the preference vectors alongside the co-fluctuation matrices in Fig. 6d,i,n. Interestingly, we observed that the extent to which a region favored unbiased random walk as a communication policy was closely linked to the mean co-fluctuation magnitude of the region. That is, two regions that prefer unbiased and diffusive communication tend to be strongly coupled to one another. In contrast, region pairs that prefer biased policies or a combination of biased and unbiased policies are not likely to exhibit correlated activity with other regions (see Fig. S3). In fact, the mean regional FC is strongly correlated with the first principal component (across top-performing models) of regional preferences for diffusive and unbiased policies (Fig. 6e,j,o).

### Exploratory analyses

In the supplement we explore, in narrow contexts, some potential follow-up extensions of this work. For instance, the framework presented here can be easily extended to include more than two policies. In Fig. S4 we examine a specific combination of policies – an unbiased random walk, shortest paths routing, and navigation (RW.wei, SP.wei, and Nav.det) and find that, as expected, the inclusion of three policies yields an improvement over all two-policy models. However, the preference patterns are not identical to any of the related two-policy models (those that pair at least two of the three policies together).

Throughout the main text we reported that high-performing models tended to combine unbiased and biased policies. What happens if we gradually make the biased policies more unbiased by incorporating some stochasticity in their routing mechanisms? One way to do this is to relax shortest path policies so that, rather than considering communication along a single shortest path, communication takes place along an ensemble comprised of the *k*-shortest paths between a source and target node [12]. Here, we calculate the *k* = 128 shortest paths between all pairs of nodes. Doing so impacts the target-dependent transition matrices for shortest paths so that, instead of every source node having a single outgoing connection, they can have up to *k* (although note that it may be the case that the first step in all *k* shortest paths is through the same node, in which case the single outgoing connection is preserved). We find that increasing the number of shortest paths, i.e. *k >* 1, leads to reductions in performance (Fig. S5a). We also tested whether there was an effect of varying which of the *k* shortest paths we retained. Again, we found clear decreases in performance whenever we considered shortest paths beyond *k* = 1 Fig. S5b). Finally, rather than fixing *k* to be equal for all pairs of regions, we thresholded shortest paths to retain those whose costs were below a particular cutoff, allowing for variability in the number of shortest paths between pairs of regions. Consequently, if two regions are connected by many paths, all of relative low cost, they may have more shortest paths between them than two regions connected by only a few low-cost paths. Here, we found that the correlation with static FC was relative poor compared to the other models, but followed an inverted u-shaped curve (Fig. S5c).

Collectively, the results of these two exploratory analyses suggest, first, that the incorporation of additional policies should be performed carefully. If the third policy performs similar to either of the first two, it may not yield much improvements in performance. Secondly, the inclusion of additional shortest paths (which introduces stochasticity in terms of where a particle gets delivered) can be viewed as making the centralized routing mechanism more diffusion-like. Our findings are directly in line with our previous analyses, which suggested that combining two unbiased diffusion processes yields poor results.

## DISCUSSION

### Joint communication policies for predicting FC

One of the longstanding questions in network neuroscience is how the brain’s anatomical connections shape patterns of correlated activity [35, 36]. While it is generally accepted that, over short timescales (on the order of minutes) FC reflects the outcome of communication events unfolding over the largely static SC, the precise policies that the brain uses to signal between regions remains unclear [4].

This question has been investigated from a number of angles, ranging from biophysically-plausible dynamical systems models [37] to amechanistic explanatory models [38, 39]. Recently, a third category of model has generated a great deal of interest. Communication models use simple processes, e.g. shortest paths routing or random walks, to explore how the structure of anatomical networks facilitates or inhibits the communication between pairs of brain regions. Communication models have a number of advantages over existing methods, including their computational tractability and ease of interpretation.

Nonetheless, communication models in their current form have a number of limitations. First, adjudicating between models has proven challenging. In most applications, model fitness (and by extension, plausibility) is assessed by comparing the capacity of communication between pairs of brain regions with the corresponding FC magnitude. Although a reasonable means of assessing fitness, in practice, most models achieve comparable performance, making it difficult to compare models to one another. Additionally, other studies select policies *a priori* and therefore never actually compare different models.

Second, communication models are typically applied at the whole-brain level. That is, it is assumed that every pair of regions communicates using an identical policy. Here, at least, some progress has been made, as several recent studies have begun to assess the regional heterogeneity in optimal communication policies [26–28]. Nonetheless, modeling communication policies locally rather than globally is uncommon.

Finally, it has proven challenging to incorporate multiple policies within the same model. Presently, the state of the art for modeling SC-FC coupling is to estimate communication matrices for several different policies and to include these matrices as predictors in a multi-linear model [13, 25]. Note that, while the models include contributions from each policy, the predictor matrix for each policy is estimated independently so that the communication policies never truly interact with one another and their combined effect remains unknown.

Here, we address these limitations by modeling shortest paths routing and greedy navigation as a target-dependent biased random walk. This allows us to seamlessly merge communication processes, traditionally viewed as distinct, with familiar diffusion and unbiased random walk dynamics. We parameterize these joint communication models regionally, so that individual nodes in the network can flexibly select the policy that, across all possible target regions, maximizes the correspondence of stationary distributions with static FC.

We show that, in combining different policies into the same model, we achieve correlations with static FC that outperform traditional communication models. On one hand, this observation directly challenges existing literature that has, historically, focused on individual communication policies in isolation. Importantly, we find that the tested models are tiered, such that when we combine unbiased diffusive policies with heavily biased and targeted policies, the outcome (in terms of the stationary distribution of random walkers) is correlated with static FC. In contrast, when we pair unbiased with unbiased or diffusive with diffusive, we observed a clear drop-off in performance.

These observations are analogous to other proposed organizational and functional principles of brain networks. For instance, it has long been argued that brain networks architectural features that balance functional integration and segregation, e.g. efficient processing paths *versus* modules [40–42]. In the present study we consider unbiased and decentralized processes and pit them against biased, target-dependent, and centralized processes – our findings suggest that a balance of these two opposed policies is necessary to reproduce, even approximately, the observed correlation structure of fMRI BOLD data, i.e. static FC. Tuning communication policies too far in either direction, e.g. pairing two centralized or decentralized policies together, leads to a stark reduction in performance.

This observation has important implications for our understanding of interregional communication. With few exceptions [23, 27], models of communication processes – including biophysical and explanatory models – assume that brain dynamics or communication policies are uniform across cortex. That is, the rules of communication are homogeneous across all neural elements. Our results suggest that this assumption leads to long-term outcomes, i.e. stationary distributions, that are inconsistent with observed coupling patterns. Incorporating multiple diverging policies, on the other hand, yields massive improvements in model fit.

On one hand, the improvements we see here could be explained on the basis of increased model complexity; the joint-policy models have more parameters than traditional communication measures and therefore *should* perform better. On the other hand, we observe considerable heterogeneity across joint-policy models. Specifically, those that pair biased with biased and unbiased with unbiased policies lead to poor performance, despite the fact that they also include additional parameters. These observations suggest that increases in model complexity alone are not sufficient for the model to improve its performance. Moreover, the results also depend heavily on the underlying network topology. Holding model complexity constant but rewiring the structural connectivity data away from its empirical organization generally leads to decrements in model performance.

Notably, we also find a clear system-level preference for certain classes of policies. Specifically, we find that sensorimotor and attentional systems exhibit preferences for unbiased diffusive processes, whereas higher-order association cortices prefer target-specific and biased policies. On one hand, these preferences may reflect underlying features of the anatomical and functional networks themselves. One of the hallmarks of both is that sensorimotor systems appear highly modular [19, 27] and a diffusive process initiated from within those modules could efficiently spread throughout the entire module, thereby supporting high levels of system-specific coupling. On the other hand, the preference for targeted and biased policies in association cortices may reflect the more complex computations thought to take place in these areas [43, 44] or increasingly diverse connectional fingerprints [45].

A possible criticism of this work is that the communication models are too stylized and sufficiently divorced from the underlying biophysics [22]. We note, first, that we model communication this way by design. In general, biophysical models are not analytically tractable and incur prohibitive computational costs, making exhaustive parameter searches and model-fitting nearly impossible [46, 47]. In contrast, simulating communication using Markov chains requires relatively little computational overhead. Second, we note that, microscopically, the role of biophysics may dominate, but at coarser scales, the collective behavior of the microscopic interactions can be reasonably modeled as simpler processes. As a relevant example, consider neural mass models, which include biophysical parameters for conductances, ion concentrations, and propagation velocities (among others) and, as output, generate synthetic membrane potential time series at sub-millisecond resolution. Despite the effort to maintain a relatively high level of neurobiological plausibility, recent work has shown that the correlation of activity simulated by these models is closely recapitulated by the dynamics of a pure diffusive process evolving over the connectome [48–50]. Nonetheless, there remains a clear need to ground communication models in neurobiological reality. The framework proposed here, which relaxes the necessity that interregional communication be homogeneous across the entire brain, represents a small step in that direction.

### Neurocognitive implications

One of the interesting observations was that regions with a preference for diffusive policies, i.e. the unbiased random walks, tend to be strongly functionally connected to one another. In contrast, regions that favor shortest paths routing or navigation exhibit weaker functional connections. These observations lead us to make two speculative claims. On one hand, these findings suggest that the key communication mechanism underlying functional coupling of regions to one another is diffusive in nature and *not* their ability to communicate selectively along a shortest path. This observation is in line with previous studies demonstrating that shortest paths matrices are relatively poor predictors of static resting FC compared to matrices based on diffusion or other decentralized processes, e.g. communicability [13–15, 25, 26].

These observations lead us to our second speculative claim. One of the more consistent findings in the analysis of task-evoked FC is that, compared to rest, it is characterized by a loss of system-level segregation [42, 51, 52]. The interpretation is that during a task specific subsets of brain regions or systems that are uncoupled and functionally autonomous at rest need to communicate with one another in order to satisfy task constraints. These inter-system connectivity increases, along with reduced intra-system connection weights [53], result in an overall reduction to system-level segregation.

Here, we find that low-levels of coupling reflect regional preferences for biased and centralized communication process, e.g. shortest paths routing or navigation. Therefore, a possible explanation for task-related increases and decreases in FC is a shift in communication preferences. Specifically, we predict that pairs of regions whose FC increases during tasks may be shifting their preferences towards unbiased policies, whereas region that exhibit task-related reductions in FC tend to favor biased policies.

More generally, our multi-policy framework makes it possible to examine changes in regional communication policies both dynamically, by applying this approach to time-varying network data, and across tasks, by fitting the multi-policy models to task FC rather than rest. Notably, we find that the organization of time-varying cofluctuation matrices can be well-approximated by multi-policy models. Because the summation across time of co-fluctuation matrices is precisely the static FC matrix, and because the co-fluctuation cluster centroids are predicted by different communication policies, we might speculate that the static FC matrix is, in fact, the superposition of many distinct communication events, each of which was driven by a different communication policy [34]. The topic of time-varying structure-function coupling is, in general, understudied and has not been fully explored using communication models (but see [28, 54]).

### Future directions

A key contribution of this present work is the mapping of communication processes, e.g. shortest paths routing and greedy navigation, onto target-dependent and biased random walks, making it possible to incorporate these processes into a multi-policy Markov chain. Here, we explore a representative but far from exhaustive set of models. Future work should be directed to investigate other possible communication policies or reparameterizations of those explored here. For instance, to estimate shortest paths, we used a reciprocal weight-to-cost transformation, which is equivalent to 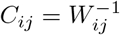. This transformation can be parameterized as 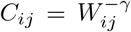, which can dramatically change the shortest paths structure of a network and, potentially, impact the performance of a joint communication model [55].

Relatedly, this work focuses exclusively on human network data at the group level. Because the aim, here, is to prove that this general framework has some broad utility, this is sufficient. Future studies, however, should extend this work from the group level to individual subjects and to network data reconstructed using other methodologies, e.g. tract-tracing for anatomical networks [56–58].

While our work provides a framework for integrating multiple interacting communication policies within the same model, like traditional uni-policy models, it does not offer an explanation for *where* the policy preferences originate and *why* particular regions and system prefer the policies they do (other than the fact that this distribution of preferences leads to stronger correlations between predicted and observed FC). In general, a first principles theory of interregional communication is presently absent; new modeling approaches begin to fill this gap, but fully addressing this area likely requires cooperation between currently separate sub-fields within neuroscience.

### Limitations

This study has several notable limitations. One of the inadvertent issues it faces is the use of correlated fMRI BOLD data (functional connectivity) as a target to which the models will be fit. Although there is a broad correspondence between the BOLD signal and population-level activity [59], the recorded signal is indirect measure of that activity and its correlation structure can be influenced by non-neural sources [60]. So while we follow other studies in selecting to use fMRI BOLD FC as the target of our model, there exist alternative possibilities. Future studies should investigate alternatives, e.g. interregional transcriptomic similarity [61] and other non-MRI markers.

In this study we focus on anatomical networks reconstructed from diffusion MRI using tractography. Although we take care in ensuring that this procedure results in connectomes that are as unbiased as possible, networks reconstructed in this way have known limitations [62–64]. Reconstructing fiber tracts with high levels of accuracy and fidelity remains an ongoing challenge for the network neuroscience community [65–68].

## MATERIALS AND METHODS

### Connectome dataset

In this study we examined network-based models of interregional communication. We carried out these comparisons using diffusion spectrum MRI data parcellated networks at a single organizational scale (*N* = 114 nodes). Here, we describe those processing steps in greater detail.

#### MRI acquisition

70 healthy participants (age 28.8±9.1yo, 43 males) were scanned on a 3T scanner with a 32-channel head coil (Magnetom TrioTim, Magnetom Prisma, Siemens Medical, Germany). The session included (1) a magnetization-prepared rapid acquisition gradient echo (MPRAGE) sequence (1×1×1.2 mm resolution, 240×257×160 voxels; TR = 2300 ms, TE = 2.98 ms, TI = 900 ms); (2) a diffusion spectrum imaging (DSI) sequence (2.2×2.2×3 mm resolution; 96×96×34 voxels; TR = 6100 ms, TE = 144 ms; q4half acquisition with maximum b-value 8000 s/mm^2^, one b0 volume). Informed written consent was in accordance with institutional guidelines and the protocol was approved by the Ethics Committee of Clinical Research of the Faculty of Biology and Medicine, University of Lausanne, Switzerland.

#### MRI preprocessing

The individual connection matrices where computed using the open aggregation software Connectome Mapper (https://connectome-mapper-3.readthedocs.io/en/latest/) [69] which calls different tools at different processing steps using the parameters described in the sequelae.

MPRAGE volumes were segmented into white matter, grey matter and cerebrospinal fluid using FreeSurfer software version 5.0.0 [70]. Cortical volumes were segmented into five progressively finer parcellations, with 68, 114, 219, 448 and 1000 approximately equally-sized parcels [71]. Here, we analyze the 114-parcel division. DSI data were reconstructed following the protocol described by Wedeen and colleagues [72], thus estimating an orientation distribution function (ODF) in each voxel. Up to three main streamline orientations were idenntified in each voxel as the maxima of the ODF (DiffusionToolkit software, http://www.trackvis.org/dtk).

Structural connectivity matrices were estimated for individual participants using deterministic streamline tractography on reconstructed DSI data, initiating 32 streamline propagations per diffusion direction per white matter voxel [73]. The MPRAGE and the brain parcellation were linearly registered to the subject diffusion space (b0) using a boundary-based cost function (FreeSurfer software) [74]. For each starting point, streamlines were grown in two opposite directions with a fixed step size equal to 1 mm. As the streamline entered new voxels, growth contributed along the ODF maximum direction that produced the least curvature. Streamlines were terminated if changes in direction were greater than 60 degrees/mm. Tractography completed when both ends of the streamline left the white matter mask. Structural connectivity between pairs of parcels was estimated in terms of streamline density, defined as the number of streamlines between two parcels normalized by the mean length of the streamlines and the mean surface area of the parcels.

### Receptor density maps

Normative receptor density maps were take from [29]. In this study, the authors aggregated PET images (1239 participants in total) from 19 neurotransmitter receptors and transporters, reflecting nine distinct systems: dopamine, norepinephrine, serotonin, acetylcholine, glutamate, GABA, histamine, cannabinoid, and opioid. Parcellated data were accessed from https://github.com/justinehansen/hansen_receptors-1/tree/main/data/PET_parcellated/scale060 and further processed by excluding subcortical regions, averaging map types across different data sources, and z-scoring the maps across cortical regions. This procedure results in a matrix of [114×19]. We treated rows of this matrix, which represent individual cortical regions/parcels, as feature vectors and computed their pairwise similarity as a bivariate correlation.

We used these data to weight edges in an unbiased random walk. To do this, we needed to remap the correlation coefficients from the interval [−1, 1] to [0, 1]. To do this, we used the transform *s* = (*r* + 1)*/*2, where *r* is the bivariate correlation coefficient and *s* is the remapped value. We calculate this measure for all pairs of regions and reweight existing structural connections with the corresponding receptor map similarity value.

### Modeling joint communication policies

#### Unbiased random walks on networks

Here, we present a framework for jointly modeling pairs of communication policies. Specifically, we present a method for mapping certain classes of deterministic communication policies onto a random walk. In general, a random walk on a network can be modeled as a discrete time Markov chain in which a walker or particle traverses a network, moving from node *i* to *j* following *i*’s outgoing connections. In an unbiased random walk, the probability of transitioning from node *i* to *j* is given by:

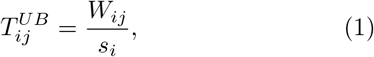

where *W*_*ij*_ is the weight of connection between nodes *i* and *j* and *s*_*i*_ = Σ_*j*_ *W*_*ij*_ is the weighted degree of node *i*. Rather than model trajectories of individual particles performing a random walk, we can equivalently model their probabilistic flow. Let *x*_*i*_(*t*) correspond to the density of particles or random walkers concentrated on node *i* at time *t*. We can calculate the density of walkers on node *i* at time *t* + 1 as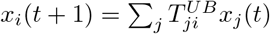.

We can also estimate the density of random walkers as *t*→ ∞, a quantity known as the stationary distribution. This value can be estimated spectrally based on the eigenvectors of *T*, or using the power method, whereby *T* is raised to some large power. For a large enough power, the rows of the exponentiated matrix are identical and are equal to the stationary distribution.

#### Biased random walks on networks

However, random walks can be biased, so that certain transitions are selectively suppressed (made less likely) or enhanced (made more likely). Here, we show that for some communication policies, e.g. shortest paths routing and greedy navigation, we can model these processes as a biased random walk.

As an example, consider shortest paths communication, in which a signal is selectively delivered from source node, *s*, to a target node, *τ*, along the shortest path *p*_*s*→*t*_ = *s, i, j*, …, *k, l, τ*. For a given target node, *τ*, we can think of the *n*−1 shortest paths (one per source) as a biased random walk. Specifically, the transition probability from node *i* to *j* is 1 if, in the shortest path *p*_*i*→*t*_, node *j* is visited immediately after *i*. Otherwise, *p*_*i*→*τ*_ = 0. We also set *τ* to be an absorbing state by giving it a self-transition probability of 1. The set of shortest path transitions – or the transitions for other target-dependent and biased communication policies – can be encoded in a square transition matrix *T* (*τ*)^*B*^ whose rows sum to unity. Notably, each row contains only one nonzero entry – all other entries are zero.

Modeling shortest paths using the language of random walks opens up a number of interesting possibilities. Most notably, we can combine the biased and unbiased random walks in a joint Markov chain by introducing the global preference parameter, *π*. Here, *π* governs the probability that a particle moving over the network will choose its next step based on the biased random walk (with probability *π*) or the unbiased walk (with probability 1 − *π*). We can write this joint random walk as:

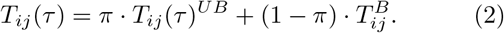

In this expression, the parameter *π* acts globally. That is, brain regions have equal preference for one or the other policy. However, we could extend this model to include regionally specific biases by allowing *π* to vary for each region, i.e. *π*_*i*_ ≠ *π*_*j*_. In this case, we would modify the above expression to read:

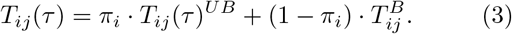

Intuitively, this expression corresponds to a random walk over the network where, when a walker arrives at node *i*, it chooses (with probability *π*_*i*_) to deliver the node to whichever of its neighbors is next on the shortest path to the target node, *τ*. With probability 1 −*π*_*i*_, the node delivers the particle to one of its neighbors in an unbiased way. Note that the neighbor on the shortest path to the target is included in both policies. We can then calculate the stationary distribution of this Markov chain in the usual ways.

Before moving to the next section, we draw the reader’s attention to a few important points. Note that the joint transition matrix is dependent upon the target node, *τ*. This is because, in general, the shortest path from node *i* to *τ* is not the same as *i* to *τ* ^′^ ≠ *τ*. Consequently, different target nodes have different transition matrices. If we consider all possible targets, we end up with *N* matrices in total.

#### Linking joint communication policies with FC

In the previous subsection, we described a framework for jointly modeling pairs of communication processes as random walks (Markov chains). We motivated this procedure by considering a joint model in which the two policies were an unbiased random walk and shortest paths routing. The proposed framework, however, is very general and can accommodate other policies, e.g. navigation. In fact, the framework could even accommodate more than two policies. Irrespective of which policies are being considered, how do we link the outcomes of this simulation to FC?

In short, our strategy for doing so is to compare FC to the stationary distribution of random walkers, i.e. under a given joint policy, the spatial pattern of where walkers end up after many steps. More specifically, we do the following. For a given target node, *τ*, a set of regionally defined preferences Π ={ *π*_1_, …, *π*_*N*_}, and a pair of policies *u* and *v*, we calculate the corresponding transition probabilities, 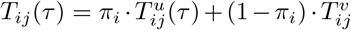. Doing so for all {*i, j*} yields a full matrix, which we raise to a large power (*t* = 500 steps). Any row of the exponentiated matrix would yield an estimate of the stationary distribution (as *t*→ ∞ every row should be identical). We calculate the stationary distribution as the average over all rows to further reduce any numerical imprecision. The result of this procedure is the vector: *x*(*τ*)^*^ = [*x*(*τ*)_1_, …, *x*(*τ*)_*N*_]. We repeat this procedure for every target node, *τ*, and arrange these vectors into the columns of a *N*×*N* matrix and symmetrize it. Finally, ignoring the diagonal elements, we extract all of the upper and lower triangle weights and calculate their Spearman correlation with upper triangle elements of the static FC matrix.

#### Simulated annealing

To fit the regional preference values of a joint model to observed FC, we use a simulated annealing algorithm. In this algorithm, we start with a set of policies and a random set of preferences. We then estimate stationary distributions for all targets and compare the results with the FC matrix using the procedure outlined above. We then select a node at random and add a small amount of Gaussian noise to its preference (clipping the value to the interval [0, 1]). We run the simulation again and estimate a new correlation between the stationary distribution and FC. If this number represents an improvement, then we retain the solution. If no, we retain the solution with probability equal to 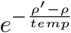 where *ρ*^′^ is the new correlation and *ρ* is the previous. Here, *temp* is a temperature parameter that gets reduced from an initial value of 1 according to the following equation *temp*(*step* + 1) = 0.999 · *temp*(*step*). This annealing schedule ensures that for large temperature values a wide range of solutions can be explored; as the temperature decreases, the algorithm is less likely to explore sub-optimal solutions, retaining on those that improve the correlation. We perform this procedure for 20000 steps and repeat the algorithm 10 times with different random preferences.

### Communication models

Previously, we described the procedure for jointly modeling pairs of communication policies. In previous work, however, we and others have considered models that include only a single communication policy applied uniformly over the entire brain. In this section, we describe (briefly) some of these measures. Note that this particular list is taken directly from [26] and is not intended to be exhaustive.

#### Flow graphs

A flow graph is a transformation of a network’s (possibly sparse) connectivity matrix, *W*_*ij*_, into a fully-weighted matrix in which the dynamics of a Markov process are embedded into edge weights [75]. Flow graphs have been applied in neuroscience for the purposes of community detection [76] and for tracking the propagation of tau deposition [77]. For a continuous time random walk with dynamics 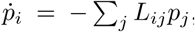, the corresponding flow graph is given by *W* ^l^(*t*)_*ij*_ = (*e*^−*tL*^)_*ij*_ *s*_*j*_. In these expressions, the matrix *L* is the normalized Laplacian whose elements are given by *L*_*ij*_ = *D*−*W/s*, where *s*_*i*_ = Σ_*j*_ *W*_*ij*_ is a node’s degree or weighted degree and *D* is the degree diagonal matrix (a square matrix with the elements of *s* along its diagonal). The variable *p*_*i*_ represents the probability of finding a random walker on vertex *i*.

The element *W* ^′^ (*t*)_*ij*_ represents the probabilistic flow of random walkers between nodes *i* and *j* at time *t*. Here, we generated flow graphs using both binary and weighted structural connectivity matrices at evaluated them at different Markov times, *t*. Specifically, we focused on *t* = 1, 2.5, 5, and 10. We refer to these variables as *fgbin*- or *fgwei* - followed by *Markov time, t*.

#### Navigation

The aim of many networks is to move something from one point in the network to another in as few steps as possible, i.e. to take advantage of shortest paths. However, doing so requires requires full knowledge of a network’s shortest path structure, which may not be a realistic assumption, especially for naturally-occurring biological systems like brains. However, it may be the case that simple routing strategies – rules or heuristics for how to move from one node to another – can sometimes uncover optimal or near-optimal shortest paths. One such routing rule is, given a target node *τ*, to always move towards the node nearest the target in some metric space, e.g. Euclidean space.

Recently, this navigation approach was applied to brain networks [9]. This study defined two novel measures based on navigation of connectome data. First, they defined the number of hops in the shortest path uncovered by the navigation process. We refer to this variable as *nav-num*. Note that for some node pairs, the navigation procedure leads to a dead end or a cycle – in which case the number of hops is listed as ∞. For the completed paths, the authors also defined their total length in metric space (in this case Euclidean distance). We refer to this variable as *nav-ms* and, like *nav-num*, impute incomplete paths with values of ∞.

#### Communicability

Communicability [78] is a weighted sum of walks of all lengths between pairs of nodes. For a binary network, it is calculated as *G* = *e*^*W*^ or 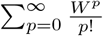. The contribution of direct links (1-step walks) is 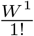, two-step walks is 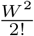, three-step is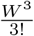, and so on. In other words, longer walks have larger denominators and, effectively, are penalized more severely. We denote this measures as *comm-bin*.

For weighted networks, we follow [79] and first normalize the weighted connectivity matrix as *W* ^′^ = *D*^−1*/*2^*WD*^−1*/*2^ where *D* is the degree diagonal matrix. As before, this normalized matrix is the exponentiated to calculate the weighted communicability *G*_*wei*_ = *e*^*W*^′. We denote this measures as *comm-wei*.

#### Matching Index

The matching index [80] is a measure of overlap between pairs of nodes based on their connectivity profiles. Suppose Γ_*i*_ = *j* : *W*_*ij*_ *>* 0 is the set of all nodes directly connected to node *i*. We can calculate the matching index between nodes *i* and *j* as 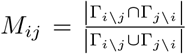. Here, Γ_*i*\*j*_ refers to the neighbors of node *i* excluding node *j*.

#### Shortest paths

In a network, each edge can be associated with a cost. For binary networks, the cost is identical for each edge; for weighted networks the cost can be obtained by a monotonic transformation of edges’ weights to length, e.g. by raising an edge’s weight to a negative power. The shortest path between a source node, *s*, and a target node, *τ*, is the sequence of edges *π*_*s*→*τ*_ = {*W*_*si*_, *W*_*ij*_, …, *W*_*kτ*_}that minimizes the sum *C*_*si*_ + *C*_*ij*_ + … + *C*_*kt*_, where *C*_*si*_ is the cost of traversing the edge linking nodes *s* and *i*.

Here, we calculated shortest paths matrices for the binary network (where the cost is identical for all existing edges) and also for a parameterized affinity-to-cost transformation evaluated at several different parameter values. Specifically, we used the following transformation: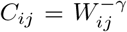. We focused on the parameter values *γ* = 0.125, 0.25, 0.5, 1.0, 2.0, and 4.0. We refer to these measures as *pl-bin* and *pl-wei-* followed by *γ value*. Note that for the multi-policy models, we only consider *γ* = 1 transformations.

#### Cosine Similarity

The cosine similarity measures the angle between two vectors, *x* = [*x*_1_, …, *x*_*P*_], and *x* = [*y*_1_, …, *y*_*P*_]. Specifically, it measures 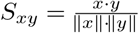. Here, we treated regions’ connectivity profiles (the row of the connectivity matrix) as vectors and computed the similarity between all pairs of regions. We repeated this procedure for both the binary (*cos-bin*) and we weighted (*cos-wei*) connectivity matrices.

#### Search Information

Search information measures the amount of information (in bits) required to traverse shortest paths in a network [25, 81]. If the shortest path between nodes *s* and *τ* is given by *π*_*s*→*τ*_ = {*s, i, j*, …, *k, l, τ*}, then the probability of taking that path is given by: *P* (*π* _*s*→*τ*_) = *p*_*si*_ × *p*_*ij*_ × … × *p*_*kl*_ × *p*_*lτ*_, where 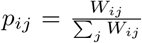. The information required to take this path, then, is *S*(*π*_*s*→*τ*_) = log_2_[*P* (*π*_*s*→*τ*_)].

Here, we calculated search information based on binary shortest paths (*si-bin*) and based on shortest paths obtained from each of the weight-to-cost transformations (*si-wei-γ value*).

#### Mean First Passage Time

The mean first passage time (MFPT) refers to the expected number of steps a random walk must evolve for a random walked starting at node *i* to end up at node *j* [82, 83]. Here, we expressed the columns as z-scores to remove nodal (column) biases and analyzed the resulting matrices for the binary (*mfpt-bin*) and weighted (*mfpt-wei*) connectivity matrices.

#### Euclidean Distance

The final predictor that we considered was the Euclidean distance between regional centers of mass (*euc*).

### Null models

We compared our results against two distinct null models. For the degree and strength preserving model, null networks were generated using the standard edge-swapping algorithm [84]. While these networks preserve nodes’ degrees exactly, they do not preserve weighted degree, i.e. strength. In general, preserving strength and degree is not straightforward. Our strategy for doing so was to first generate a binary network whose degree sequence was identical to that of the observed network. Next, we used a greedy algorithm to configure the original weights over the randomized edges such that the degree sequence was as similar as possible to that of the original network.

For the degree, strength, and geometry preserving null model, we adopted a nearly identical approach for network generation. In this case, we used a class-specific implementation of the edge-swapping algorithm, wherein we only allowed edge swaps to occur between edges of similar connection length (Euclidean distance). This constraint results in networks whose cost was similar to that of the original network.

### Edge time series and cluster analysis

Recent studies have shown that static FC can be temporally unwrapped into its framewise contributions [30, 31]. The procedure for doing so is straightforward. Note that the correlation between two variables, *x* = [*x*(1), …, *x*(*T*)] and *y* = [*y*(1), …, *y*(*T*)] can be written as:

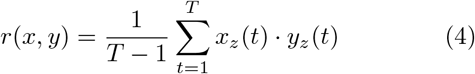

where 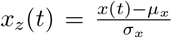 and *μ*_*x*_ and *σ*_*x*_ is the mean and standard deviation of *x*.

In our previous work, we demonstrated frames corresponding to high-amplitude co-fluctuations, i.e. “events”, corresponded to co-fluctuation patterns that were repeated across scans, were highly identifiable of individuals, and could explain large percentages of variance in the static FC [30, 32, 33]. Here, we applied an event detection algorithm to whole-brain edge time series from all subjects, extracted the corresponding co-fluctuation patterns, and clustered them using modularity maximization. Note that this procedure is identical to the one described in [34]. This algorithm resulted in three distinct clusters that were shared across many individuals. We treated the centroids of these clusters – the average across all co-fluctuation patterns assigned to each cluster– as putative network states.

---

**FIG. S1.**
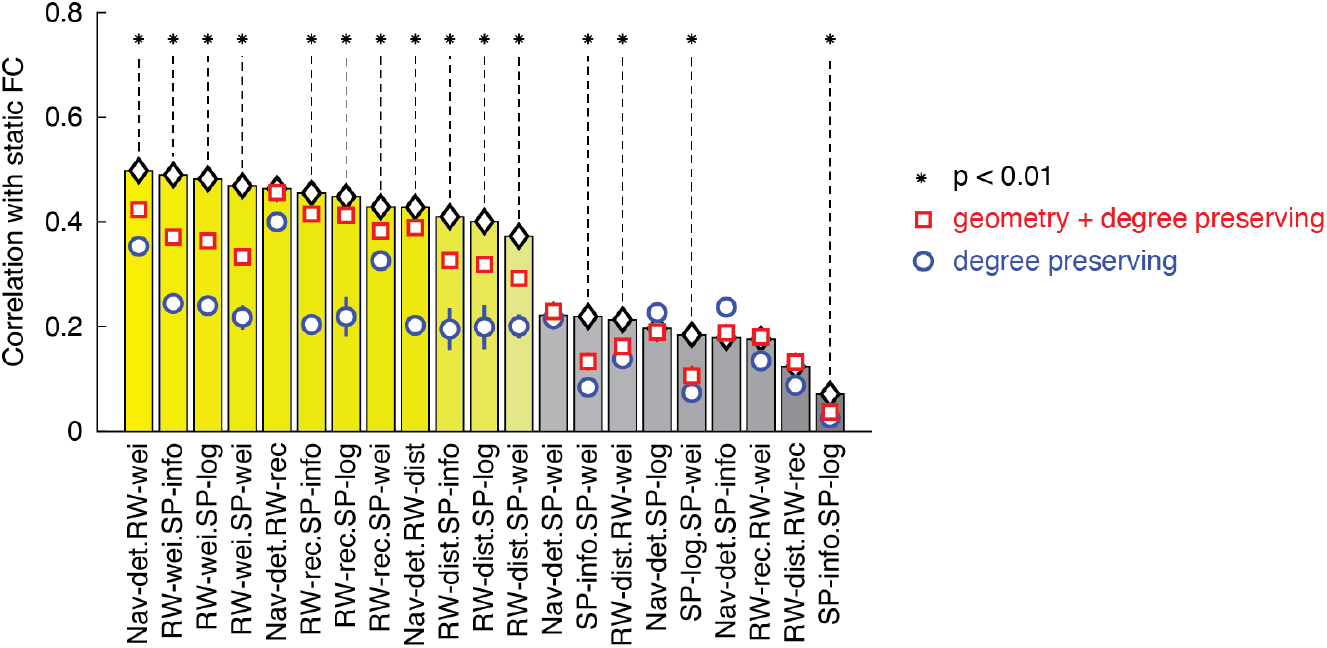
Comparing joint models fit to empirical networks to fits using randomized networks. In the main text, we report the structure-function correlation after fitting joint models to static FC. Here, we repeat the same analysis using randomized structural connectivity data. We explored two null models. Both models preserve nodes’ degrees exactly while also approximately preserving nodes’ strengths. Additionally, one of the models also preserves (approximately) the total wiring cost. In this figure, we denote the degree + strength + geometry preserving model with red squares and the degree + strength model as blue circles. Asterisks indicate that the structure-function correlation using the observed network outperformed the null networks (*p <* 0.01, uncorrected).

**FIG. S2.**
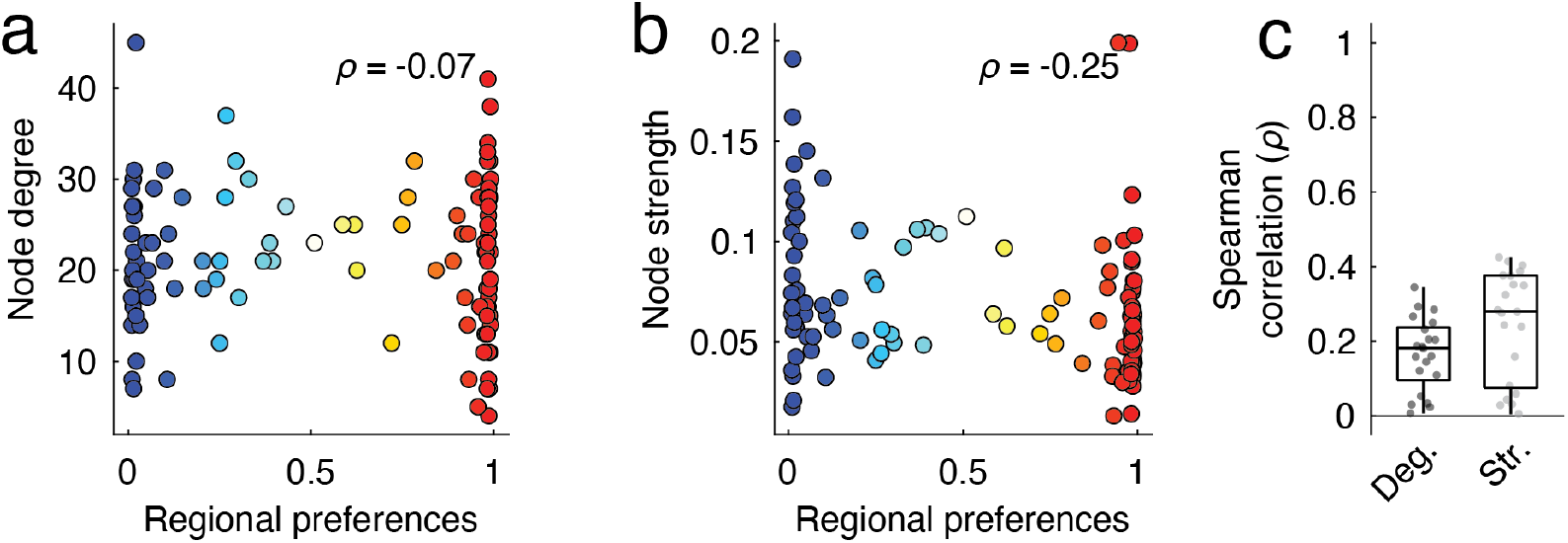
Comparing regional policy preferences with low-level features of structural connectivity. In the main text we fit multi-policy models by varying regional preferences one policy or the other. Here, we show that these policy vectors are largely uncorrelated with node degree and strength. (*a*) Correlation of regional preferences with node degree for the best-fitting model fit to static FC (Nav.det-RW.wei).

**FIG. S3.**
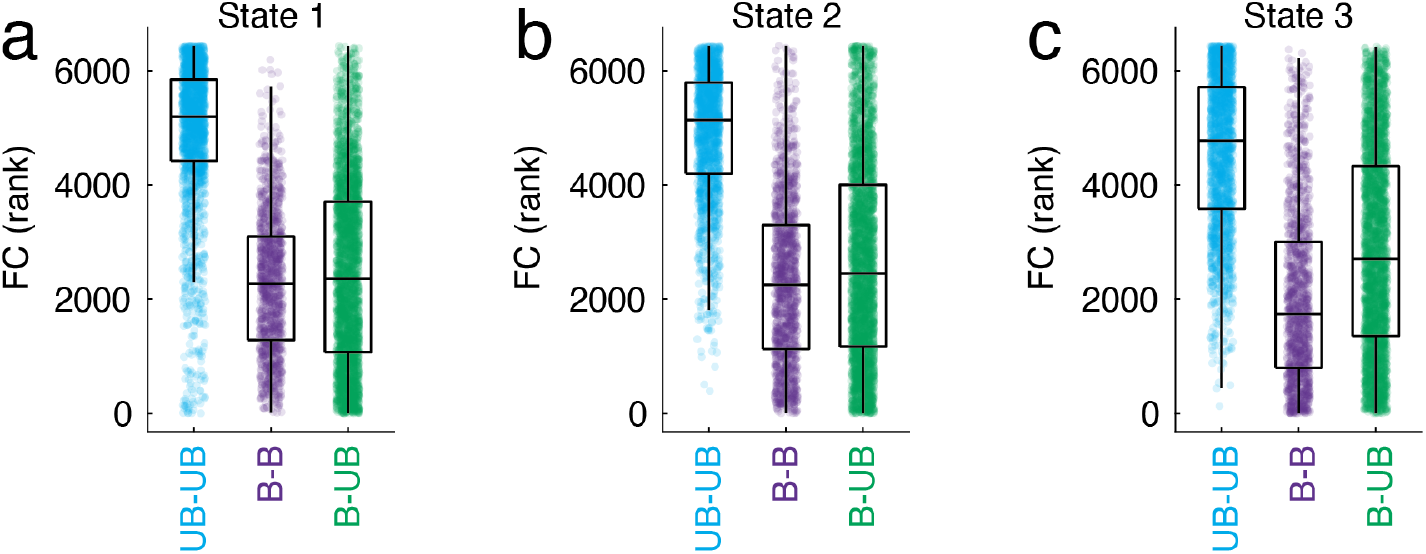
Linking dynamic fluctuation to edge categories. In the main text we described a procedure for categorizing edges into three groups. Briefly, we considered joint models in which one policy was a diffusive and unbiased random walk and another was based on a centralized and target-dependent process. Every region could have a preference for one policy or the other. We considered a region *i* to prefer the unbiased policy if its preference was *π*_*i*_ *>* 0.5. We labeled these regions as RW regions. Otherwise, we labeled the region SP. We could then classify all region pairs based on their stub nodes’ assignments. This results in three possible pairings: RW with RW, SP with SP, and RW with SP. Here, we show that the dynamic co-fluctuations between RW+RW pairs tend to be stronger than for the other two edge pairings. We show this for states 1, 2, and 3 (panels *a, b*, and *c*).

**FIG. S4.**
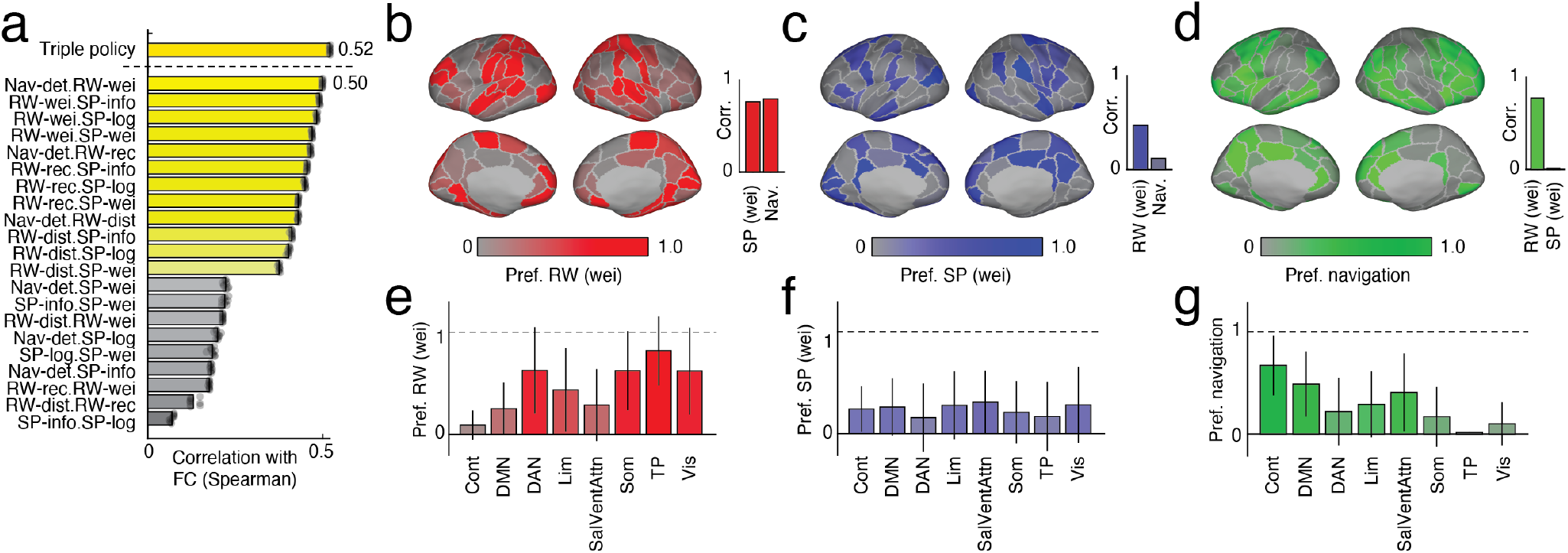
Exploratory analysis of three-policy model. In the main text we examined two-policy models. Here, we explore a three-policy model combining RW.wei, SP.wei, and Nav.det policies. (*a*) In general, we find that three-policy model significantly outperforms the best two-policy model (*ρ* = 0.52± 0.001; t-test, *p <* 0.01). In panels *b*-*g* we show regional preference vectors for the three policies. We find that the unbiased weight-based random walk favors sensorimotor, attentional, and temporoparietal systems while the navigation policy favors the control and default mode networks. In contrast, the shortest paths policy has no clear preference at the system level.

**FIG. S5.**
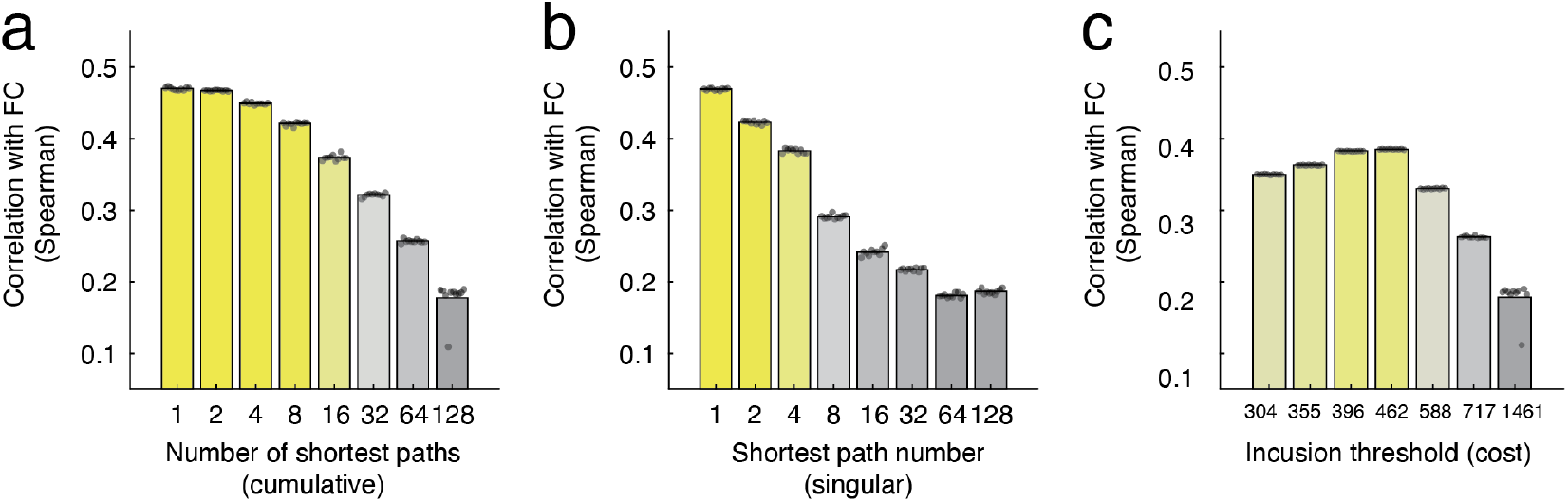
Exploratory analysis using *k*-shortest paths. In the main text we examined communication *via* shortest paths. Here, rather than considering only one shortest path, we assess the effect of communication *via* the *k*-shortest paths. Specifically, we considered two scenarios. First, rather than forcing a node to pass a particle to the target node along its shortest path, we allow it to pass the particle along any of its *k* shortest paths, selecting where to deliver the node probabilistically. In this way, the addition of paths makes the shortest path mechanism more diffusion-like. We show the results of this analysis in panel *a*. Next, rather than use the shortest overall path, we forced routing to occur along the *k*-th path, in descending order of cost (panel *b*). Finally, we include all paths of cost *c* or less.

## Notes

### Competing Interest Statement

The authors have declared no competing interest.

